# Personality profiling may help select better cleaner fish for sea-lice control in salmon farming

**DOI:** 10.1101/2021.05.21.444956

**Authors:** Benjamin Alexander Whittaker, Sofia Consuegra, Carlos Garcia de Leaniz

## Abstract

Lumpfish (*Cyclopterus lumpus*) are increasingly being used as cleaner fish to control parasitic sea-lice in salmon farming, but cleaning rates are very variable and not all individuals eat sea-lice, which increases the risk of emaciation and has ethical and practical implications. Selecting good cleaners is a priority to make the industry more sustainable, but there is little information on what behaviours make cleaner fish effective under a commercial setting. We examined variation in lumpfish personalities according to the five-factor personality model that takes into account differences in activity, anxiety (shelter use, thigmotaxis), aggression, sociality, and boldness (neophobia). We then quantified how variation in lumpfish personalities influenced interactions with naïve Atlantic salmon (*Salmo salar*), without the confounding effects of variation in sea-lice loads. Variation in activity, sociality, aggression and neophobia, but not in anxiety, was repeatable, which is consistent with a heritable basis. Neophilic, non-aggressive lumpfish spent more time inspecting salmon than neophobic and aggressive individuals, but salmon fled in the presence of the most active and social individuals, suggesting there may be an optimal cleaner fish personality amenable to artificial selection. The personality screening protocols developed in this study could inform a more efficient use of cleaner fish in salmon farming and reduce the number of individuals required to control sea-lice.

## Introduction

Various species of wrasse and lumpfish (*Cyclopterus lumpus*) are increasingly being used as facultative cleaner fish to remove parasitic sea-lice from farmed salmon (Powell *et al*., 2018b; Treasurer, 2018a; Treasurer, 2018b). Using facultative cleaner fish for pest control offers many advantages over chemical methods (Powell *et al*., 2018b), but relies on cleaner fish engaging in parasite removal (delousing) under a commercial setting. However, lumpfish do not generally feed on sea-lice, or act as cleaner fish, in the wild (Davenport, 1985; Powell *et al*., 2018a), which poses a challenge for optimizing their use in salmon farming. Lumpfish are opportunistic generalist feeders that will graze on sea-lice in salmon cages, especially when alternate food sources are limited (Imsland *et al*., 2015a; Imsland *et al*., 2016a). Typically only 13-38% of lumpfish consume sea-lice in salmon cages (Eliasen *et al*., 2018; Imsland *et al*., 2015a; Imsland *et al*., 2014; Imsland *et al*., 2016b), the majority feed on salmon pellets or graze on plankton (Johannesen *et al*., 2018), particularly during the summer (Eliasen *et al*., 2018). This has attracted criticism from regulatory agencies and animal welfare organisations, who point out that many cleaner fish die of malnutrition and suffer from low welfare because they are being used for a task they do not naturally do (European Union Reference Laboratory for Fish Diseases, 2016; Stranden, 2020). A call has been made for the use of cleaner fish to stop until their welfare can be guaranteed and the sources of mortality are addressed (Compassion in World Farming, 2018; OneKind, 2018). There is, therefore, a need to select individuals that readily engage in cleaning behaviour and consume sea-lice, as this would increase delousing rates with fewer cleaner fish, improve welfare, and reduce the risk of emaciation (Gutierrez Rabadan *et al*., 2020).

Lumpfish can be effective cleaners (Imsland *et al*., 2014; Imsland *et al*., 2018) and reduce sea-lice loads on salmon by as much as 93% under the right conditions (Imsland *et al*., 2016b), but seemingly this is because certain individuals do most of the cleaning. Individual variation in cleaning behaviour is common among facultative cleaner fish, and some individuals may attend clients and engage in cleaning, while others ignore them (Morado *et al*., 2019). Understanding such variation is important because cleaning behaviour appears to be inherited (Imsland *et al*., 2016b), and the ability to screen individuals with a predisposition for cleaning may facilitate a selective breeding program (Powell *et al*., 2018b). However, there is a paucity of robust data on individual variation in cleaning behaviour and little information on lumpfish-salmon interactions (Overton *et al*., 2020), which has so far hampered progress towards the domestication of lumpfish as cleaner fish in aquaculture.

Domestication requires animals to adapt to life in captivity (Price, 1999), to express some behaviours and to suppress others (Belyaev, 1969; Jensen, 2014). Juvenile lumpfish aggregate in clumps and are sit and wait feeders (Powell *et al*., 2018a; Powell *et al*., 2018b). They are also poor swimmers (Hvas *et al*., 2018), have a low aerobic scope (Killen *et al*., 2007b), and use their suction cup to attach to shelters to conserve energy (Killen *et al*., 2007a). They are not well adapted to pursue prey, and do not normally interact with other fish species. To function as cleaner fish, lumpfish may need to overcome the fear of approaching a much larger “client” fish, change their social behaviour, and become more exploratory, active and inquisitive. More specifically, delousing salmon requires lumpfish to take four successive steps: (1) leave the safety of being in a group and/or in a shelter, (2) approach a much larger and faster swimming fish (salmon), (3) identify clients that are infested with sea-lice, and (4) remove sea-lice from the skin of a fast moving target. This sequence of events requires a particular skillset that not all individuals may possess, which might explain why only a small proportion of lumpfish clean parasitic sea-lice in salmon farms. We therefore hypothesized that only those individuals that have a bold personality, show exploratory behaviour, or are more active and willing to take risks may interact with salmon. Identifying those individuals might be possible through personality profiling *sensu* McCann (1992).

Personality profiling has been used to understand variation in personalities in humans (Costa Jr & McCrae, 1990; Neal *et al*., 2012; Van Dijk *et al*., 2017), but increasingly also in non-humans (Gosling & John, 1999), where it is referred as temperament or stress coping styles (Dingemanse *et al*., 2010; Piersma & Drent, 2003). Personality traits are repeatable in many taxa, including fish (Castanheira *et al*., 2017; Elias *et al*., 2018; Vargas *et al*., 2018), which opens possibilities for artificial selection and the domestication of cleaner fish in aquaculture. We therefore tested if personality profiling using the five-factor model of non-human animal personality (Gosling & John, 1999) could be used to identify lumpfish better suited to behave as cleaner fish in salmon farms. We also tested if behaviours relevant to delousing were repeatable, as only repeatable behaviours can be inherited and modified by artificial selection during animal domestication (Belyaev, 1979).

## Materials and Methods

We used a four-step approach to assess the effects of lumpfish personality on lumpfish-salmon interactions. We first screened individually tagged lumpfish for five dimensions of animal personality (activity, sociality, anxiety, aggression, and boldness; (Gosling & John, 1999) in the absence of salmon. We then retested fish after ~1 month to assess the repeatability of behaviour along each dimension, and generated personality scores from Principal Component Analysis to account for correlated behaviours. Finally, we examined whether lumpfish personality scores predicted lumpfish-salmon interactions.

### Source and rearing of experimental fish

Lumpfish eggs were collected from wild adults caught in Iceland and the English Channel during the winter of 2016/2017, representing two genetically distinct populations (Whittaker *et al*., 2018). Juveniles were reared for one year (mean weight = 114.8 ±9.7 g SE) under recirculation aquaculture conditions and a sample allocated at random to eight 1500L tanks (1.4 m diameter, 0.9 m depth; 33 fish/tank; initial biomass = 75.37 ±30.99 g/m^3^) and individually marked with PIT tags (7 x 1.35 mm, Loligo) in March 2017. Water temperature was maintained at 11-13C, salinity at 28-32ppt, and photoperiod at 12D:12L; fish were fed twice per day (Amber Neptune Skretting, UK) at 2% tank biomass, as recommended for the species (Powell *et al*., 2018b).

### Personality profiling

We filmed 38 individually tagged lumpfish of two origins (Iceland, n = 16; England, n = 22) over four consecutive phases, each lasting 10 minutes, using CCTV (1080p camera, Sannce, Hong Kong). In phase 1 (neutral), a single lumpfish was introduced into one of two white rectangular test arenas (L120 cm x W55 cm x D25 cm) and was left to acclimatise for 10 mins without any additional stimulus. The test arenas were divided into three equal zones using tape to facilitate video analysis, were each fitted with an air stone to maintain dissolved oxygen at 100% saturation and were surrounded by a black tarpaulin screen to minimise disruption. In phase 2 (shelter) a black smooth PVC panel was introduced to one end of the test arena to provide a refuge, as lumpfish seek smooth, dark substrates (Imsland *et al*., 2015b). In phase 3 (mirror), a mirror was added to the arena, opposite to the shelter. Finally, in phase 4 (novel), a novel object was placed at the centre of the arena (**Figure 1**), a yellow golf ball in the first trial and a green Lego brick in the second repeat to prevent habituation. The four screening phases were conducted in the same order for every fish. Water was replaced between fish to avoid a build-up of stress hormones.

**Figure 1.**
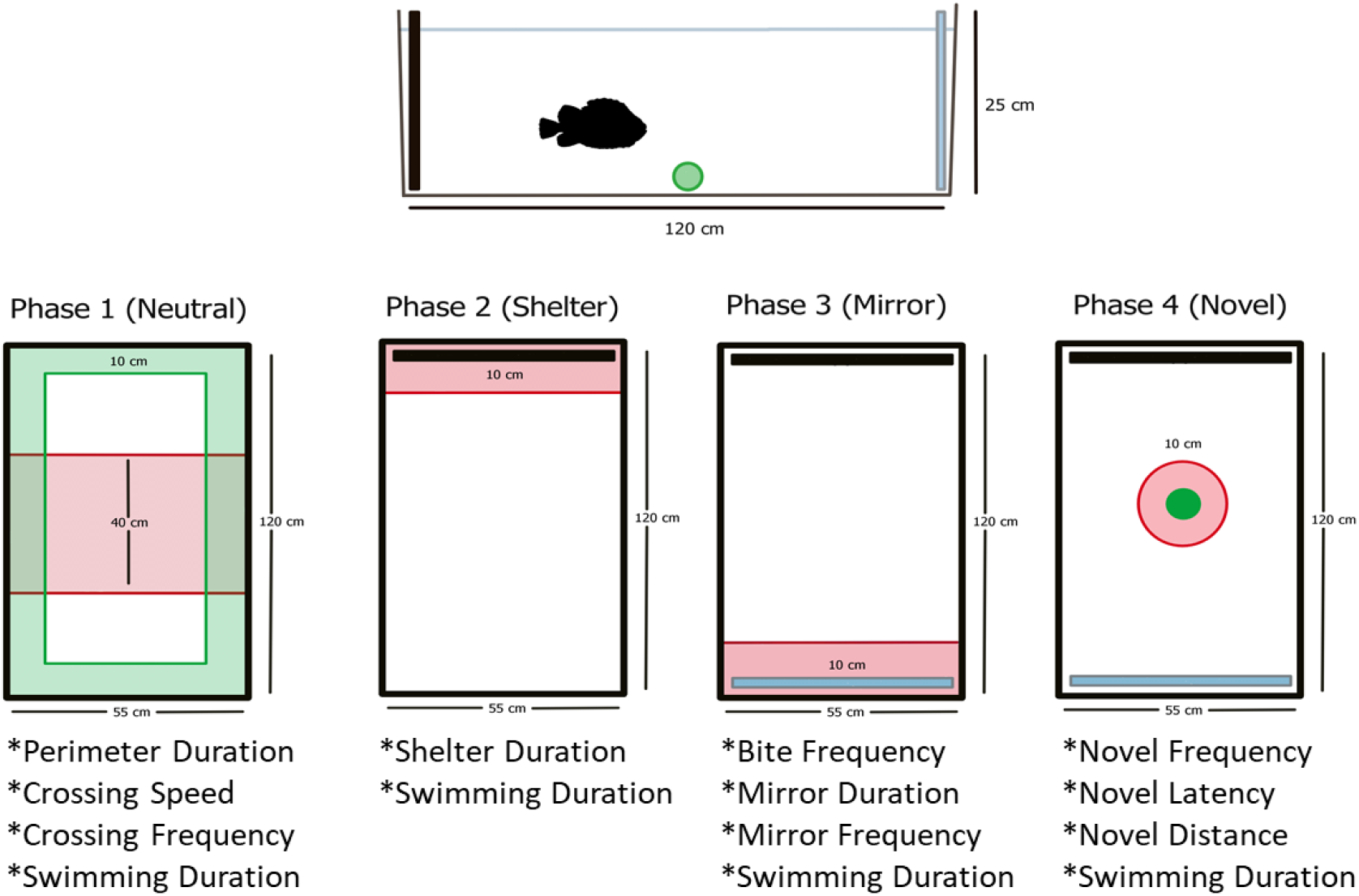
Experimental design showing the four experimental phases, each lasting 10 minutes to asses variation in lumpfish personalities. In the neutral phase (phase 1), a perimeter was superimposed over the video footage to assess time spent near the walls (thigmotaxis) in an open field test and three equal sections were marked to measure crossing frequency and swimming speed. In the shelter phase (phase 2), the time spent within 10 cm of a black panel was recorded. In the mirror phase (phase 3), the time and frequency of agonistic and non-agonistic approaches to the mirror were recorded. In the novel phase (phase 4), the latency, frequency of approaches within 10 cm and distance to the novel object were recorded.

Behaviours were scored from video recordings using BORIS version 7.75 (Friard & Gamba, 2016) by one person who did not have knowledge of the identity of the fish or the trial conditions. We examined two metrics of activity in phase 1 (neutral): crossing speed across the central portion of the test arena (cm/s), and number of crosses between the three arena sections. We also measured activity by recording the total time spent swimming (s) throughout all four phases. Anxiety was assessed by recording the total time spent within 10cm of the tank walls (thigmotaxis) during phase 1 (neutral) and also the total time spent within 10cm of the black panel in phase 2 (shelter), as both are established methods of assessing anxiety in fish (Cianca *et al*., 2013; Godwin *et al*., 2012). To score aggression, we counted the number of agonistic actions (nips and charges) directed towards the mirror image (Rodriguez-Barreto *et al*., 2019) during phase 3 (mirror). The number of approaches to the mirror that did not result in body contact, as well as the time spent within close proximity (<10 cm) of the mirror during this phase were used as measures of sociality (Cattelan *et al*., 2017). As a measure of boldness, we counted the number of times individuals were within 10 cm of the novel object phase 4 (novel), the time latency to first approach the novel object, and average distance (cm) from novel object at one minute intervals, as these are measures of neophobia that map well into the boldness-shyness continuum (Champneys *et al*., 2018). Collectively, these metrics capture variation in five dimensions of fish personality and have been associated with the expression of particular genes in the fish brain (Champneys *et al*., 2018; Rodriguez-Barreto *et al*., 2019). We also measured body mass (g) and calculated specific growth rate (SGR%,(Lugert *et al*., 2016) between the first and second repeated trials to account for the influence of feeding motivation on behaviour.

### Quantifying lumpfish-salmon interactions

Atlantic salmon (*Salmo salar*) post-smolts (n = 20, mean mass = 175.6 ± 37.6g SE) were obtained from a commercial farm in Scotland in July 2017 and quarantined for one month in two 1500 litre recirculating aquaculture tanks (1.4m diameter, 0.9m depth), at a density of 10 salmon per tank (initial biomass = 117.04 ±25.06 g/m^3^ SE). In August 2017, the same tagged lumpfish that had previously been screened for personality traits (n = 38) were introduced singly into one of the two salmon tanks and recorded via CCTV for 20 minutes each. An ethogram based on previous studies on cleaner fish (Bshary & Côté, 2008; Horton, 2011) was constructed from the following three behaviours: time spent visually assessing salmon, often from multiple angles (“inspection, s”), time spent following salmon (“pursuit, s”), and number of times salmon fled when they were approached by lumpfish (“flee”, no/min). Salmon were free of sea-lice to ensure that interactions with lumpfish were unaffected by variation in sea-lice loads, thereby ensuring consistency between replicated trials and between individuals. This would not have been possible if sea-lice loads had decreased as a result of lumpfish delousing, as this might have changed the behaviour of both species. All lumpfish were naive to salmon and salmon were also naive to lumpfish initially, but they were exposed to multiple lumpfish over the course of the study.

### Repeatability of behaviours

R version 4.0. (R Core Team, 2020) was used for all statistical analyses. To asses the repeatability (*R*) of behaviours we used mixed models to identify significant behavioural covariates using the *lme4* package (Bates *et al*., 2014). We employed linear mixed models (LMM) in the case of continuous behavioural metrics (e.g. duration of events, swimming speed and distance to novel object), and generalized linear mixed-models (GLMM) in the case of frequency of events (e.g. number of crosses across the test arena, and approaches to the mirror or the novel object). We included sex, stock origin (Iceland, England), specific growth rate (SGR, %), and time of filming (days elapsed) as fixed factors, and test tank, home tank, and fish ID as random factors to account for repeated sampling and to control for grouping and potential non-independence of results. For each behavioural metric we identified minimal adequate models (Crawley, 2013) based on changes on AICc using the *dredge* function in the *MuMIn* package (Barton & Barton, 2015), and used the *anova* command and the Likelihood-Ratio Test (LRT) to compare these to null models without predictors. Model assumptions were checked by inspection of diagnostic plots with *sjPlot* (Lüdecke, 2020), and if these were not met, the significance of results was compared with robust estimates using the *robustlmm* package (Koller, 2016). The most plausible models were refitted by Restricted Maximum Likelihood (REML) to derive corrected probability values.

We used the *rptR* package (Stoffel *et al*., 2017) to calculate adjusted repeatabilities (*R*; (Nakagawa & Schielzeth, 2010; Schuster *et al*., 2017), using fish ID (n = 38) and repeat (n = 2) as grouping variables, and any significant covariates identified above; 95% confidence intervals were calculated via 1,000 bootstraps, using a gaussian or a poison link, depending on the nature of the response variables. Behaviour was considered repeatable if the lower 95% CI exceeded zero (Neumann *et al*., 2013) and repeatability was deemed low (*R* < 0.2), moderate (*R* = 0.2-0.4), or strong (*R* > 0.4), as in other studies of animal personalities (Bohn *et al*., 2017). Non-repeatable behaviours were excluded from analysis and repeatable behaviours were averaged to obtain a single value per fish.

### Personality scores and effects on lumpfish-salmon interactions

Four repeatable behaviours identified from the personality profiling (crossing frequency, approaches to mirror, charges at mirror, and approaches to novel object were converted to frequency rates (No/min) and used to generate personality scores via Principal Component Analysis to account for correlated behaviours and lack of independence (Gosling & John, 1999; Wilson *et al*., 2014). PC scores were rescaled from 0 to 1 using the *scales* package (Wickham *et al*., 2020) to facilitate comparisons and ensure equal weighting. To examine the effect of personality on lumpfish-salmon interactions we modelled the time lumpfish spent inspecting and pursuing salmon, as well as the salmon fleeing rate, as a function of lumpfish personality scores while statistically controlling for tank effects, stock origin, sex, growth rate, and time elapsing (to account for the cumulative exposure of salmon to lumpfish). We modelled each interaction separately via univariate LMMs and also together via MANOVA, and derived minimal adequate models via model simplification as described above for the repeatability study.

### Ethics declaration and approval for animal experiments

This study adhered to the ARRIVE guidelines. All experimental procedures were approved by Swansea University, Animal Welfare Review Body, permit IP1617-27.

## Results

### Repeatability of behaviours

Crossing frequency was influenced by sex (parameter estimate (male) = 0.594 ±0.222, *z* = 2.675, *P* = 0.007) and specific growth rate (parameter estimate = 0.525 ±0.195, *z* = 2.688, *P* = 0.007), as females and faster growing fish were more active and crossed the test arena more frequently. Aggression decreased the second time the fish were tested against their mirror image (parameter estimate day = −0.010, ±0.004, *z* = −2.352, *P* = 0.019), as did the frequency of non-aggressive mirror interactions (parameter estimate day = −0.005 ± 0.001, *z* = −3.783, *P* < 0.001). Stock origin influenced the time individuals spent close to the tank perimeter (parameter estimate (UK) = 56.18 ± 22.72, *t*74 =2.473, *P* = 0.016) and the number of approaches made to novel objects (parameter estimate (UK) = −1.611 ± 0.386, *z* =-4.171, *P* < 0.001), with fish from the English Channel spending more time close to the walls and making fewer interactions with the novel object than fish from Icelandic origin. We therefore included sex and specific growth rate to estimate the adjusted repeatability of crossing frequency, geographic origin to estimate the repeatability of time spent close to the walls and frequency of encounters with the novel object, and time between trials to estimate the repeatability of interactions with the mirror image.

Of the 11 behaviours considered, 4 were repeatable corresponding to the activity, aggression, sociality and boldness dimensions of animal personality (**Table 1**). Crossing speed and time spent swimming were strongly repeatable (R>0.4), crossing frequency and frequency of interactions with the mirror and the novel object were moderately repeatable (R = 0.2-0.4), while aggression had a low repeatability (R = 0.184). Distance to the novel object and the two measures of anxiety were not repeatable.

**Table 1.** Repeatability (R) of behaviours used to generate personality scores for lumpfish. Repeatable behaviours are highlighted in bold.

| Personality component | Behaviour | R | SE | *P* |
| --- | --- | --- | --- | --- |
| **Activity** | **Crossing speed (cm/s)** | **0.651** | **0.097** | **<0.001** |
|  | **Crossing frequency (No.)** | **0.278** | **0.127** | **0.018** |
|  | **Swimming duration (s)** | **0.422** | **0.135** | **0.004** |
| **Sociality** | **Approaches to mirror (No.)** | **0.375** | **0.118** | **<0.001** |
|  | Time close to mirror (s) | 0.000 | 0.080 | 1.000 |
| Anxiety | Time in arena perimeter (s) | 0.000 | 0.094 | 1.000 |
|  | Time in shelter (s) | 0.177 | 0.135 | 0.124 |
| **Aggression** | **Charges at mirror (No.)** | **0.184** | **0.238** | **0.005** |
| **Boldness** | **Approaches to novel object (No.)** | **0.340** | **0.186** | **0.033** |
|  | Latency to novel object (s) | 0.184 | 0.137 | 0.163 |
|  | Distance to novel object (cm) | 0.226 | 0.139 | 0.092 |

### Personality scores

Significant positive correlations were found between two of the four repeatable behaviours (**Table 2**). Activity was positively correlated with sociality, and sociality was positively correlated with aggression. The first two principal components accounted for 68% of the variation in personality. PC1 was mostly driven by sociality (loading = −0.64) and activity (loading = −0.58), while PC2 mostly measured boldness (loading = +0.94) against aggression (loading = −0.26). PC3 was not significant in any of the models considered.

**Table 2.** Correlation matrix between repeatable behaviours in lumpfish used to generate personality scores via Principal Component Analysis. Upper diagonal displays the Pearson’s correlation coefficient and lower diagonal the associated probabilities. Significant correlations are highlighted in bold.

| Personality component | Activity | Sociality | Aggression | Boldness |
| --- | --- | --- | --- | --- |
| Activity<br>(No. crosses/min) |  | **+0.440** | +0.210 | -0.046 |
| Sociality<br>(No. mirror approaches/min) | **0.006** |  | **+0.330** | +0.098 |
| Aggression<br>(No. mirror charges/min) | 0.210 | **0.044** |  | -0.094 |
| Boldness<br>(No. novel object approaches/min) | 0.780 | 0.560 | 0.580 |  |

### Lumpfish-salmon interactions

In the presence of salmon, most lumpfish engaged in visual inspections (89% of cases) and triggered one or more flight responses from salmon (89% of cases), but only 34% of lumpfish pursued salmon (**Figure 2)**. Lumpfish that spent more time inspecting salmon were also more likely to pursue them (Spearman rho = 0.550, df = 37, *P* < 0.001). Time spent pursuing salmon was not explained by personality (*F*_7,30_ = 1.066, *P* = 0.408) but lumpfish personality predicted the time spent inspecting salmon (PC2 estimate = −18.151 ± 8.861, *t*_34.385_ = −2.048, *P* = 0.048), as well as the frequency of salmon flights (PC1 estimate = −1.005 ± 0.242, *t*_33.419_ = −4.150, *P* < 0.001) while statistically controlling for significant random tank effects. Individuals that were not afraid of the novel object during the initial behavioural profiling (neophilic) subsequently spent more time inspecting salmon than neophobic fish, as did the least aggressive lumpfish (PC2 - **Figure 3A**). Salmon were more likely to flee in the presence of the most active and social individuals (PC1 - **Figure 3B**).

**Figure 2.**
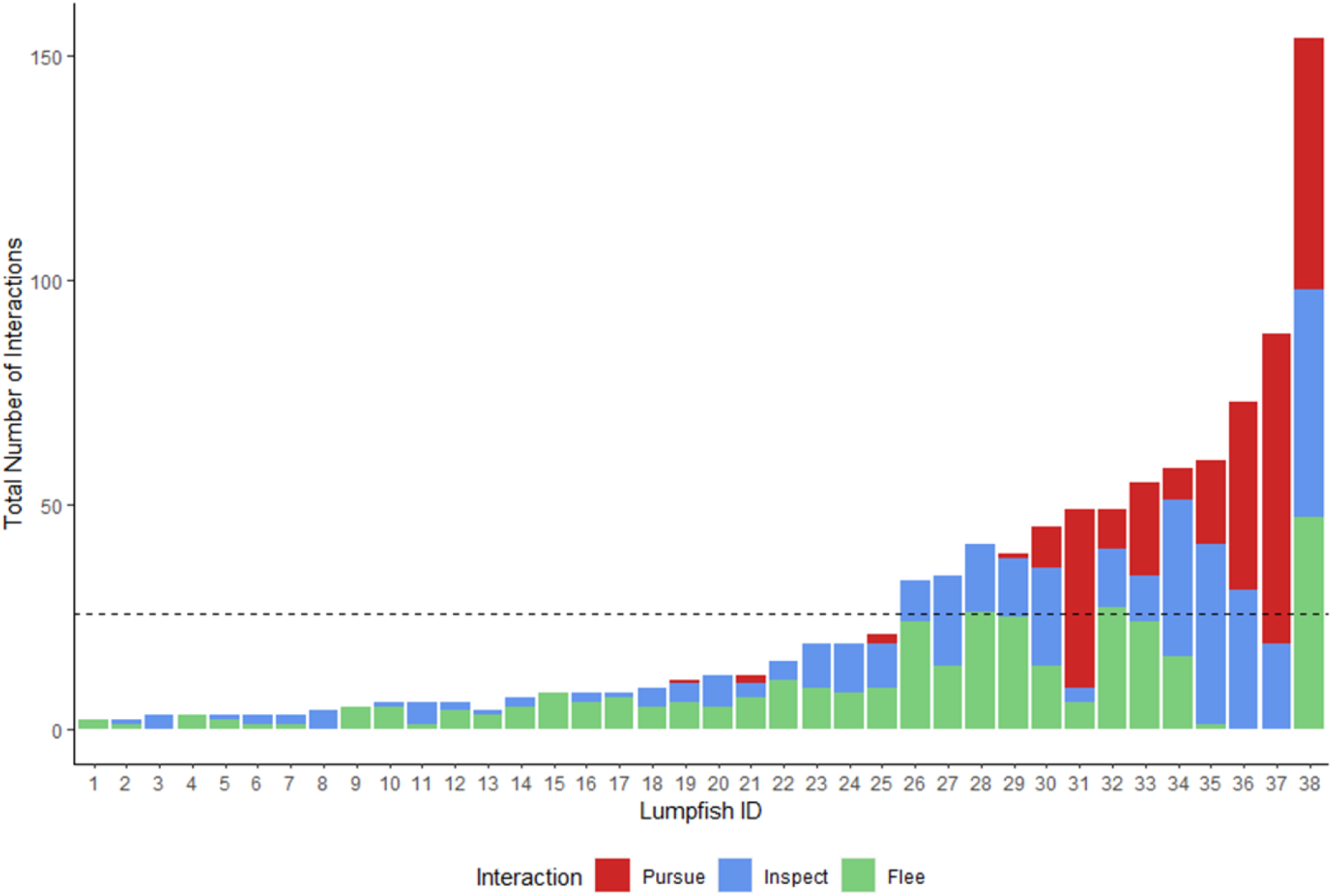
Interactions observed between single lumpfish and 10 post-salmon smolts during 20-minute observation sessions (n = 38). Dotted line shows the mean number of interactions across all observations.

**Figure 3.**
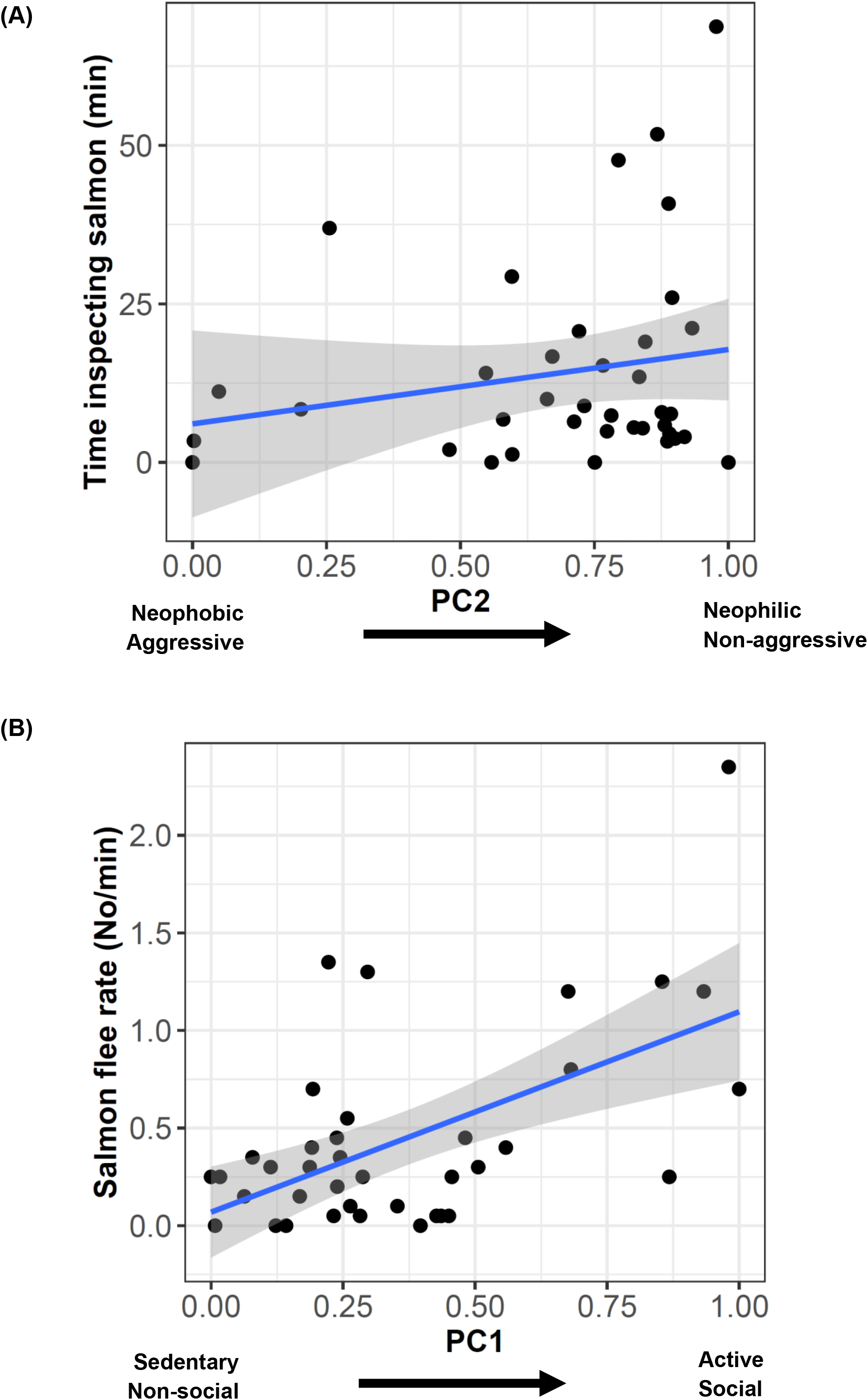
Relationship between significant lumpfish personality scores (PC1, PC2) derived from behavioural profiling and (A) time spent by lumpfish inspecting salmon, and (B) frequency of salmon flights in response to lumpfish approaches.

MANOVA indicated that lumpfish-salmon interactions were predicted by personality (PC1 Pillai = 0.422, *F*_3,33_= 8.047, *P* < 0.001) and growth rate (Pillai = 0.332, *F*_3,33_= 5.480, *P* =0.004), and provided a significantly better fit than a null model (*F*_6,68_= 5.398, *P* < 0.001).

## Discussion

Our study shows that personality profiling can be used to predict how lumpfish interact with salmon and, possibly, whether they will make good cleaner fish. In particular, we show that bold, neophilic lumpfish were more likely to inspect salmon, while the most active and social individuals were more likely to cause salmon to flee. As these behaviours were repeatable, and therefore likely heritable (Dohm, 2002; Lynch & Walsh, 1998), this suggests that artificial selection could be used to select better cleaner fish through domestication (Powell *et al*., 2018b). This is consistent with results that show that delousing behaviour is parentally controlled (Imsland *et al*., 2016b) and thus likely inherited. However, of the 37% of lumpfish that pursued salmon in our study, only 5% did so without causing salmon to flee. The challenge for domesticating lumpfish as cleaner fish, hence, might be how to select individuals that are bold enough to approach salmon in search of sea-lice, but not so aggressively as to cause salmon to flee.

The distribution of lumpfish-salmon interactions was highly skewed. The majority of lumpfish had limited interactions with salmon, but a few individuals interacted much more than the rest. For example, 7 individuals accounted for 60% of all the inspections, and just three individuals accounted for 60% of all the pursuits (Figure 2) with only 37% of them pursuing salmon. This is consistent with results from diet analysis that indicate that only ~30% of lumpfish consume sea-lice in salmon cages (Eliasen *et al*., 2018; Imsland *et al*., 2015a; Imsland *et al*., 2014; Imsland *et al*., 2016b), which our results suggest might be due to variation in animal personalities. Boldness is a significant predictor of interactions in dedicated cleaner fish (Dunkley *et al*., 2019; Wilson *et al*., 2014), and it was repeatable in our study, suggesting it could targeted by artificial selection. Likewise, activity, exploratory behaviour and aggression were also repeatable, and several studies have shown that these traits are inherited in several fish species (Ariyomo *et al*., 2013; Bakker, 1986; Chervet *et al*., 2011; Magnhagen, 2012; Sutrisno *et al*., 2011).

Breeding programs could establish pedigree lines from lumpfish families with proven delousing ability in sea-cages (Imsland *et al*., 2016b) and select for stocks with desirable behaviours. For example, fish from Icelandic origin in our study spent significantly more time interacting with the novel object than fish from the English Channel, suggesting there may be genetic differences in risk-taking behaviour that could be improved through selective breeding. There is no information on the heritability of any behavioural traits in lumpfish, but the estimated repeatabilities for swimming duration (R = 0.422) and frequency of approaches to the novel object (R = 0.340) could be used as upper values for narrow sense heritability (*h*^2^) of pursuing and inspection behaviours, respectively (Dohm, 2002; Lynch & Walsh, 1998). Using these values to estimate the response to artificial selection suggests that selecting the top 10% most active and boldest lumpfish for breeding could increase the time lumpfish spend inspecting and pursuing salmon 1.4-3.5 times within one generation, respectively. In this sense, protocols could be developed to screen lumpfish personalities in hatcheries, and exclude from subsequent rearing and deployment those individuals less likely to approach salmon. Currently, many lumpfish do not engage in cleaning behaviour, and selecting for behaviours that make good cleaners could drastically reduce the number of lumpfish required on farms, reducing costs and improving welfare (Garcia de Leaniz *et al*., 2021).

The selection of “elite” lines of cleaner fish that are particularly bold and active, but not overly aggressive, could be done in different ways. For example, one way boldness could be selected is by targeting lumpfish with high metabolic rate (Hvas *et al*., 2018) and low cortisol response (Gutierrez Rabadan *et al*., 2021), or more efficiently, by targeting genes associated with boldness and aggression (Rodriguez-Barreto *et al*., 2019). Metabolic rate can be used to predict boldness in some fish species (Killen *et al*., 2012) and can also be used to predict foraging mode (active vs passive) in lumpfish (Killen *et al*., 2007a). This is important, as presumably only active lumpfish that swim to pursue their prey (as opposite to passively foraging while clinging) will make efficient cleaners.

One reason why some farmed lumpfish do not survive the salmon production cycle in sea cages is because they fail to feed and adapt to a novel environment (Imsland *et al*., 2020; Imsland *et al*., 2019b). Failure of hatchery-reared lumpfish to adapt to the sea-cage environment could be due to phenotypic mismatch, as seen in other species (Stringwell *et al*., 2014), but steps can be taken to reduce maladaptation. Fish personality is shaped by both intrinsic and extrinsic drivers (Brown *et al*., 2011), most notably through early experience (Dingemanse *et al*., 2009; Huntingford & Garcia Garcia de Leaniz, 1997), and one way maladaptation could be reduced is by manipulating the rearing environment through environmental and social enrichment. Environmental enrichment can modify risk-taking behaviours in fish very rapidly (Roberts *et al*., 2011; Roberts *et al*., 2014) and this could perhaps be used to suppress fear and enhance cleaning behaviour in lumpfish. For example, feeding sea-lice and live prey to lumpfish prior to cage deployment was found to promote subsequent delousing behaviour (Imsland *et al*., 2019a), and previous exposure to salmon may also reduce stress and improve cohabitation (Staven *et al*., 2019). In general, intensive fish farming seems to modify fish behaviours in predictable ways, by altering patterns of gene expression in the fish brain that can be targeted by selective breeding (Rodriguez-Barreto *et al*., 2019). Social enrichment prior to deployment in sea cages could be used to modify behaviours and facilitate adaptation of lumpfish to sea cages. For example, living in socially impoverished habitats decreases cleaning efficacy in wrasse (Wismer *et al*., 2014), and the same could happen in lumpfish.

Cleaning behaviour could also be improved through size selection and manipulation of growth rate. We found that faster growing lumpfish interacted more with salmon than slower growing fish, suggesting that selecting for fast growing lumpfish might also select for more active, bolder lumpfish which might perform better as cleaner fish. Lumpfish hatcheries typically supply relatively small juveniles to sea-cages (approx. 20g), as this is thought to be an optimal size for cleaning behaviour (Imsland *et al*., 2016c). However, more recent studies indicate that body size accounts for little variation in delousing rates, and that smaller lumpfish are more likely to have empty stomachs (Eliasen *et al*., 2018). Larger cleaner wrasse also interact more with salmon (Whittaker *et al*., 2021) and a re-assessment of the optimal size for lumpfish deployment warrants further study.

In summary, our results indicate that only a small proportion of lumpfish interact with salmon in ways that are conductive to cleaning behaviour. As some personality traits were found to be repeatable, this opens the possibility for artificially selecting cleaner fish with desired behaviours, a process that our study suggests could be rapid. Failure to feed on sea-lice is a major cause of emaciation and mortality among lumpfish used as cleaner in salmon farming (Gutierrez Rabadan *et al*., 2021), and behavioural profiling could be used to select better cleaner fish and make the industry more sustainable. Further studies, using pedigree analysis of variation in delousing rates could be used to estimate the heritability of personality traits and their association with sea-lice removal.

## Acknowledgements

We are grateful to Jessica Minett and Kayla Fairfield for assistance with the personality screening and to Paul Howes, Becky Stringwell and the CSAR technicians for logistic support and help with fish husbandry. The study was funded by Marine Harvest Scotland through the LUMPFISH project and the Welsh Government via the European Regional Development Fund (SMARTAQUA Operation). The funders had no role in study design, data collection and analysis, decision to publish, or preparation of the manuscript.

## Data availability

The data files for this study have been deposited in figshare https://doi.org/10.6084/m9.figshare.14626839.v1

